# Modernization, wealth and the emergence of strong alpha oscillations in the human EEG

**DOI:** 10.1101/125898

**Authors:** Dhanya Parameshwaran, Tara C. Thiagarajan

## Abstract

Oscillations in the alpha range (8-15 Hz) have been found to appear prominently in the EEG signal when people are awake with their eyes closed, and since their discovery have been considered a fundamental cerebral rhythm. While the mechanism of this oscillation continues to be debated, it has been shown to bear positive relation to memory capacity, attention and a host of other cognitive outcomes. Here we show that this feature is largely undetected in the EEG of adults without post-primary education and access to modern technologies. Furthermore, we show that the spatial extent and energy of the oscillation have wide variation, with energy ranging over a thousand fold across the breath of humanity with no centralizing mean. This represents a divergence in a fundamental functional characteristic of an organ demonstrating both that modernization has had a profound influence on brain dynamics and that a meaningful ‘average’ human brain does not exist in a dynamical sense.

## INTRODUCTION

Oscillations in the alpha range (8-15 Hz) were among the first visually observed phenomenon in the dynamics of the human brain, described by Hans Berger shortly after he performed the very first EEG recording in 1924 with the most primitive of instrumentation. These alpha oscillations dominate the EEG signal during the awake, eyes-closed state and are considered today to be a fundamental feature of human brain dynamics[1], referred to as the ‘archetypal cerebral rhythm’[2]. Easily measured today by EEG these oscillations emerge as a distinct peak in the power spectrum of the signal with an average frequency of about 10.5 Hz. Unraveling the specific mechanisms and role of this dominant rhythm of the human brain has since been of considerable interest. The peak frequency has been shown to vary across individuals[3] and increasing evidence suggests that it has broad relation to a number of cognitive capacities[1, 4]. Either the peak frequency or strength of the oscillation have been found to correlate positively to working memory[5–7], attention[8–10] and learning ability[11] and decline with increasing age[7]. In addition, lower levels of resting alpha have been found in fragile X mental retardation syndrome[12] and in patients with Alzheimer’s and amnesic mild cognitive impairment[13] as well as ADHD[14]. Finally, substantial evidence now lends to the hypothesis that resting state alpha oscillations may correspond to an inhibition of distracting information to allow sharpened internal focus and mental imagery[15].

Here we have evaluated the alpha oscillation under an eyes closed resting paradigm across a broad range of human populations as part of a larger inquiry to understand the impact of modernization on human brain dynamics. Modern civilization represents a very rapid shift in human experience with the advent of larger urban agglomerations, more extended formal education and a host of enabling technologies including electricity, telecommunications and motorized transport. Access to these experiences of modern life are largely linked to income, a factor that is increasingly divergent across humanity. This study therefore involved recording EEG from 402 adults between the ages of 21 and 65 from across 48 settlements in India ranging from remote hamlets to large cities. Participants in the study had annual incomes spanning $300 to $150,000 and, correspondingly, a wide range of access to formal education and modern technologies. We have previously shown, using the same dataset, that the complexity of the EEG waveform differs dramatically across modern and pre-modern populations, separating into two distinct distributions[16]. We asked here how the phenomenon of modern civilization might have impacted the characteristics of the alpha oscillation.

We characterized both the peak frequency and energy of the alpha oscillation in the EEG, finding surprisingly, that the alpha oscillation was largely undetected in the EEG of populations that were premodern in their experience (and correspondingly low-income). Furthermore, we found that the energy of the alpha oscillation spanned over a 1000 fold range among our sample with no centralizing mean indicating the absence of a meaningful average.

## MATERIALS AND METHODS

### Sampling, Survey and EEG Recording

Data was gathered from 402 people across 48 settlements in the State of Tamil Nadu, India including very small hamlets of only 300 people with no electricity or motorized transport as well as large cities. Participants answered a series of questions regarding their demographic, communication and mobility behavior in addition to having EEG recorded for three minutes while they were awake and seated with their eyes closed. In 20 people, 10 recording were done in separate sessions over a week to assess intra-person variability. The EEG recordings were carried out using the Emotiv EPOC device. While the device has been shown to have similar results to more expensive clinical grade systems [17–21], we nonetheless performed validations by comparing results to simultaneous recordings from the clinical grade Neuroscan device along multiple dimensions with highly correlated results[16]. Survey methodology and sampling are described in more detail in a companion paper [16].

### Extracting Peak Frequency and Energy of the Alpha oscillation

To characterize the oscillation, rather than computing just the alpha band in the power spectral density, we use a method that magnifies the oscillation peak relative to the characteristic 1/f decay of the power spectrum, enabling detection of even very weak oscillations and separating this peak from the background decay.

To do this we first computed the discrete fourier transform of the autocorrelation using a 5 second Hann window to obtain the power spectral density (PSD). To then identify the peak frequency of the oscillation we computed the discrete approximation of the second order derivative of the PSD, and found the maximum between 7.5 and 15 Hz minus 2. We have chosen to use the FFT of the autocorrelation (R) since the autocorrelation provides an intermediate step that provides a useful visualization of the presence of periodicity. We represent it with the notation S_R_ so as to distinguish from the PSD computed using other methods.

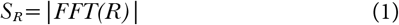

To amplify detection of the oscillation and suppress the background decay we use a second order differential of the function. This is a method commonly used in spectroscopy and other fields to amplify the detection of small signals embedded within decaying backgrounds. The higher order the differential, the greater the amplification of small signals. Here we compute *S”_R_* as the second order difference function of the FFT of the autocorrelation (Fig. 2G-I, grey line).

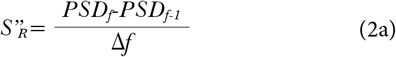

for which the numerical derivative is equivalent to

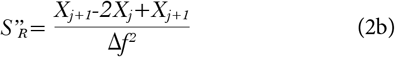

where f is frequency.

We compute the peak *F_a_* as the peak of this as

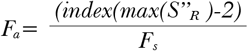

Where *F_s_* is the sampling frequency.

We then defined a measure of energy of the oscillation E_α_ as the sum of the second order derivative ± 3.75 Hz from the peak detected within the alpha band frequencies. Energy of a time series is generally computed as the integral of the second derivative of the amplitude with respect to time. We thus use a similar computation with respect to the frequency rather than time to provide the energy of the oscillation. The analog formula would be as follows.

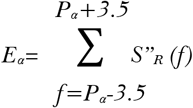

where *f* is frequency.

This method has an advantage over an integration of the power spectrum density in that it amplifies the rate of change and therefore minimizes the contribution of non-oscillatory alpha components. Nonetheless, the α energy sometimes took on a nonzero value even when there was no distinct peak detected since there was some fluctuation in the power in the α range even if an oscillation did not exist. Furthermore, it also captures information about the sharpness or fidelity of the peak frequency and not just the overall strength. We have developed a more refined method that is described in much greater detail in this paper [28].

### Statistical Tests

We made assessments of the significance of differences between groups using both the t-test as well as the non-parametric Kolmogorov-Smirnov (KS) test given that not all the distributions were normal in nature. For age and modernity where the groups were >3, we performed an ANOVA 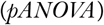 and also computed the probability of finding a similarly significant separation if the values were shuffled across participants pshuff. To do this the *P_α_* and *E_α_* values were shuffled across participants 5000 times. Pshuff represents the number of instances out of 5000 where the p-value of the ANOVA value was ≤ the pANOVA of the actual data.

## RESULTS

### Characterizing the Alpha Oscillation

The alpha oscillation or rhythm is most easily visualized in the autocorrelation of the signal. Fig. 1A shows the autocorrelation over a one second period of signal, averaged across channels, for three participants in our study. For the first participant (top), the autocorrelation is obviously dominated by an oscillation of ~10 Hz (10 cycles over the one second duration shown). For each participant we analytically determined both the peak frequency (*P_α_*) of the oscillation for each channel and the energy (*E_α_*) of the oscillation. The energy can be thought of as a function of both the amplitude and fidelity of the oscillation to a particular frequency (see Methods).

**Figure 1.**
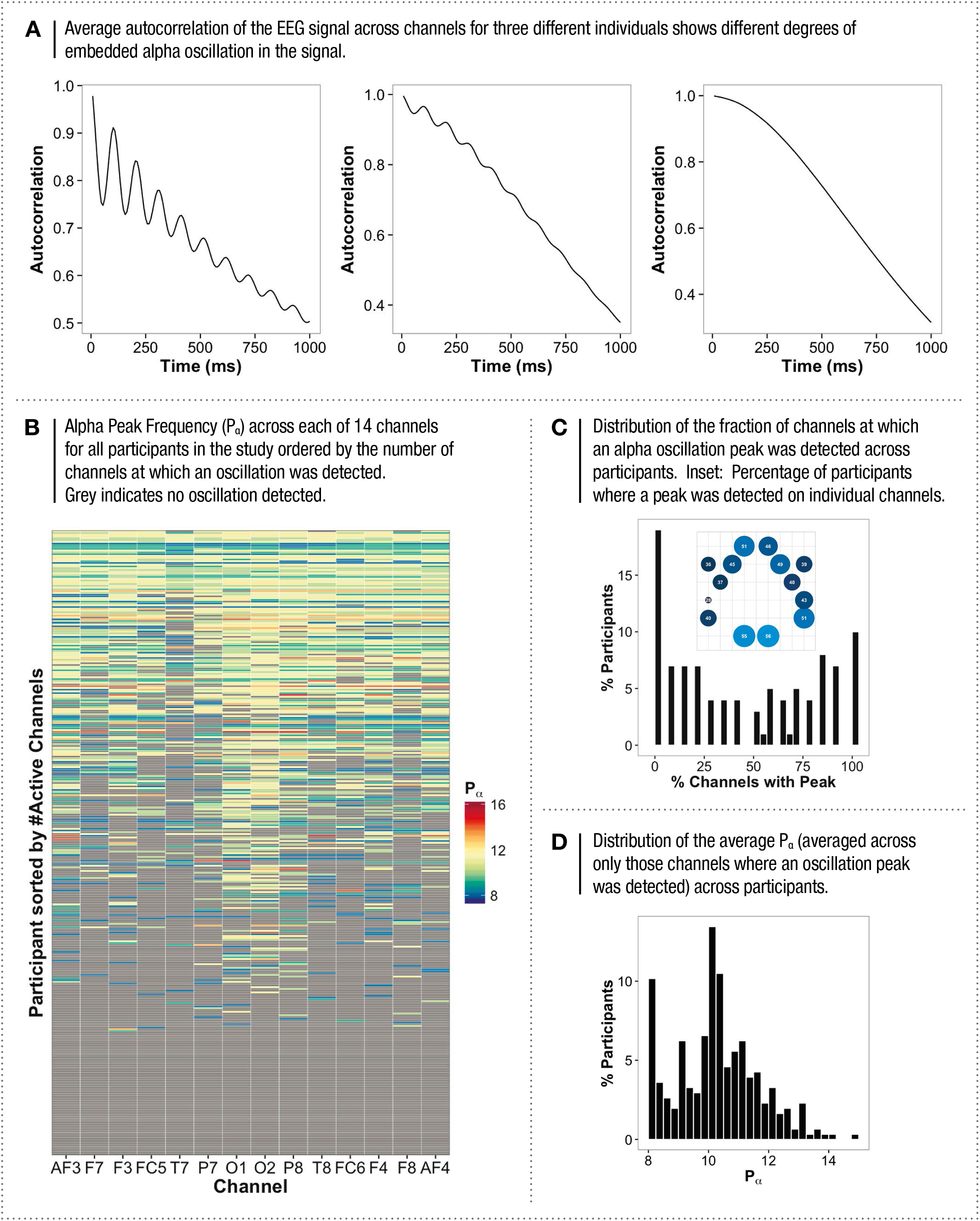
ALPHA OSCILLATION PEAK FREQUENCY (P_a_) ACROSS OUR SAMPLE

Fig 1B shows *P_α_* for each channel for each participant (each row is a participant). Surprisingly we found that 20% of the participants showed no evidence of an oscillation in any of the 14 channels recorded, which can be seen as a smooth average autocorrelation (Fig. 1A, bottom) and as grey in the matrix in 1B. Overall there was large variation in the fraction of channels where an oscillation was detected among participants spanning the spectrum from 0 to 100% (Fig. 1C). The oscillation was most consistently present in the occipital region (O1,O2 Fig. 1C inset) followed by the frontal and prefrontal areas. It was least present in the area T7 (left parietotemporal cortex).

Thus while there was wide variation, given that we did not find a complete absence of the oscillation across all ten recordings in any of the 20 individuals, this suggests that it is unlikely that some human brains do not have the mechanism to produce the alpha oscillation. Furthermore, evidence for alpha oscillations have been found in other species as well, particularly monkeys [22, 23] leading to the suggestion that it is an intrinsic property of the architecture of the mammalian brain[24]. However, it is significant that these recordings in other species have not been done with the EEG but rather with implanted electrodes directly on the brain surface recording local field potentials (LFPs) or electrocorticographs (ECOG) at significantly higher spatial resolution, well below the limits of detection of the non-invasive EEG. This suggests that this feature is not sufficiently strongly developed in some people so as to enable detection at the spatial resolution of the EEG.

Taken together our results suggest a very broad continuum rather than an absolute absence of the feature in some people.

For those channels that had a detectable oscillation, the average peak frequency across participants (computed by averaging the frequencies across those channels where it was detected) was 10.5 Hz, consistent with previous studies[3]. The standard deviation (SD) and coefficient of variation (CV%) of *P_α_* across participants were 2 Hz and 19% respectively, again closely in line with previous studies. However, clearly the presence of the oscillation varied greatly across participants, and the fraction of channels with a peak was not significantly correlated with the peak value across channels (*r*=0.11).

To get deeper insight into the variation across the population we recorded activity in ten separate instances over a one-week period in 20 people. The 20 people selected were spread across the spectrum of having the oscillation present in all channels in the initial recording to being absent in all channels. In this subset, when an oscillation was present we found an average *P_α_* in the sample of 10.62 Hz with an intra-person SD of 1.1, Hz in line with previous studies[3]. However we found a very wide intra-person variation in the number of channels where an oscillation was detected (Fig. 2A-C). The fraction of channels with a detectable peak across recordings varied across participants with individual participants having a cross-recording mean ±SD of 0.92 ±0.06 (highest) to 0.12 ±0.1 (lowest). Across the sample, the oscillation was present in at least one channel anywhere from 60% to 100% of the time (Fig. 2C). That said, the average *P_α_* was relatively stable in most individuals with a median intra-person (cross-recording) CV% of 6.05% compared to the median across individuals, which was 15.4% (Fig. 2D).

**Figure 2.**
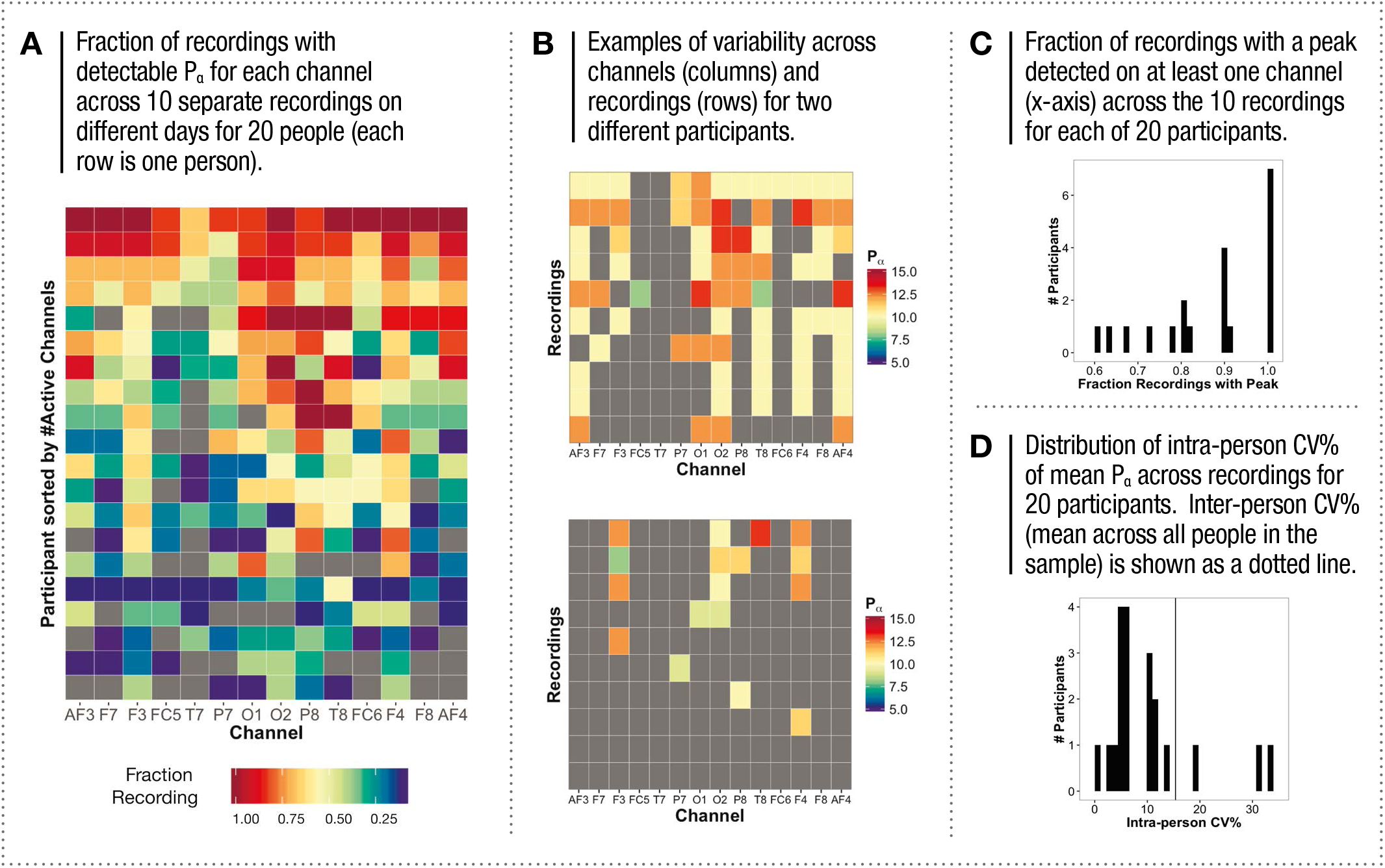
INTRA-PERSON VARIATION IN ALPHA PEAK FREQUENCY

### The Alpha Energy (E_α_)

Fig. 3A shows the energy of the alpha oscillation (*E_α_*) across channels (columns) and participants (rows) ordered as in 1B (see Methods for details of definition). Here again there was large variation both across channels and participants. While the measure is designed to amplify detection of the oscillation peak rather than residual alpha component in the spectrum, in some channels where a peak could not be detected, there was nonetheless some residual energy in the alpha band. For channels with no detectable alpha oscillation, *E_α_* was normally distributed with a mean ± SD of 21±9 (Fig. 3B). Interestingly, however, for those channels where an alpha peak was detected, the average energy across channels for all participants ranged from 10 to over 1350 in a distribution that was heavily skewed or long tailed with no centralizing characteristic (Fig. 3C). The mean value (134±185) is therefore not meaningful to this structure; the median was 67 and energy in an individual channel went as high as 5680. The median intra-person CV% for the average *E_α_* across channels was 56% compared to the median inter-person CV% of 137% (For. 3D) indicating that the differences across the population sampled far exceeded the differences from recording to recording within individuals. We also note that given that the distributions of *P_α_* and *E_α_* were fundamentally different, these two measures were not correlated (*r* = 0.09).

**Figure 3.**
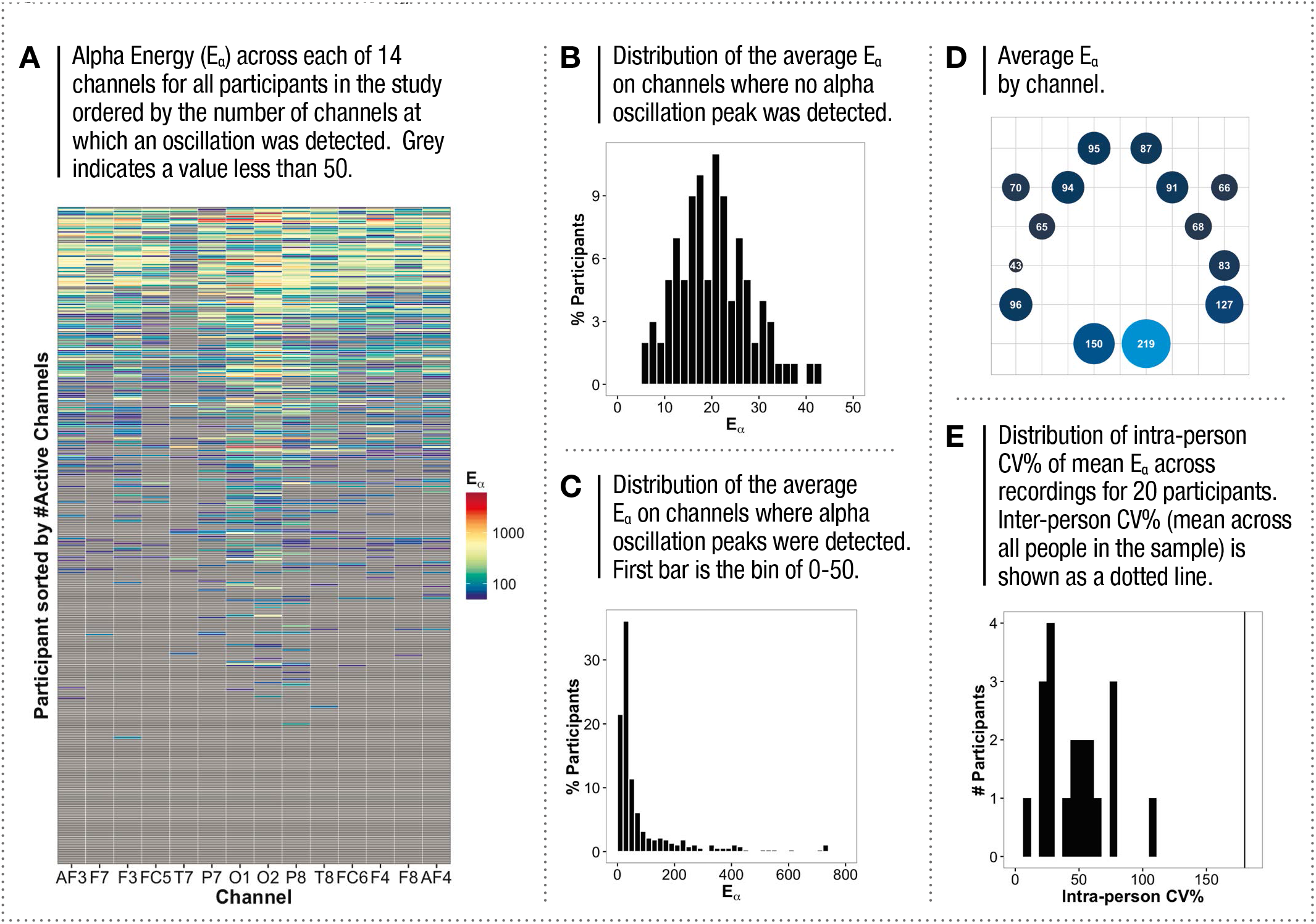
ALPHA ENERGY (E_α_) ACROSS OUR SAMPLE

Although studies have demonstrated differences in *P_α_* across regions [25], the alpha oscillation is most strongly present in the occipital region making it possible that a very strong oscillation in this region might be detected at other channels in the EEG simply due to volume conduction. To rule this out we looked at the fraction of channels at which an oscillation was detected as a function of the energy of the oscillation in channels O1 and O2 (Fig. 4A). In both cases (shown here is O2 which was stronger than O1 overall and very similar), while a very strong oscillation in O1 or O2 (>250) was overall more likely to be associated with oscillations detected on more channels, it was nonetheless clear that oscillations could often be present on all channels even when *E_α_* of the occipital region was very low. Furthermore, if detection of the oscillation at other channels arose through volume conduction, *E_α_* in O1 and O2 would be highly correlated to the *E_α_* of other regions. However, the mean occipital *E_α_* compared to the mean *E_α_* at other regions had a linear R^2^ of only 0.37 (Fig. 4B) suggesting that oscillations detected on other electrodes were largely mediated by other mechanisms. In addition, peak frequencies while highly correlated within the frontal/prefrontal and occipital regions, were not very correlated across these two regions (Fig. 4C), which would not be expected if the origin of frontal alpha were on account of volume transmission from the occipital lobe. This is consistent with various reports that suggest differences in the alpha oscillation frequency across brain regions[3, 9, 26].

**Figure 4.**
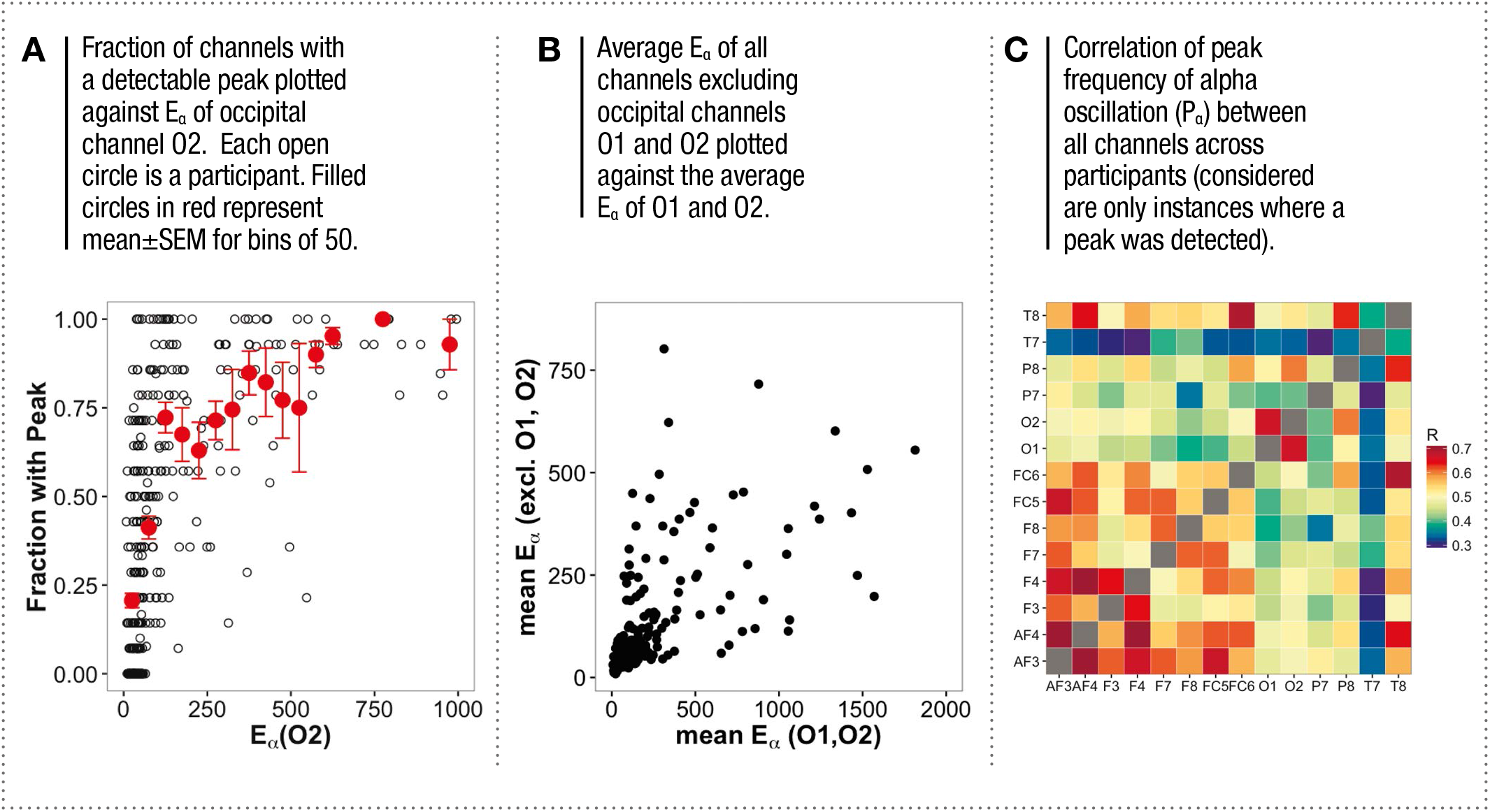
ALPHA ENERGY AND SPATIAL CHARACTERISTICS

## THE ALPHA OSCILLATION, AGE AND GENDER

We looked first at the features of the alpha oscillation as a function of gender (Fig. 5A,B). Earlier studies have reported slightly higher alpha frequency in adult females relative to males[27]. While we found a similar directional difference, the magnitude was not large enough to be significant in our dataset. *P_α_* was 10.27±0.10 for males and 10.71±0.19 for females (Fig 5A; 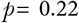 by KS-test, 0.06 by t-test). *E_α_* was 98.77±12 for males and 93.36±11.15 for females (Fig. 5B; 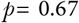 by KS-test, 0.74 by t-test).

**Figure 5.**
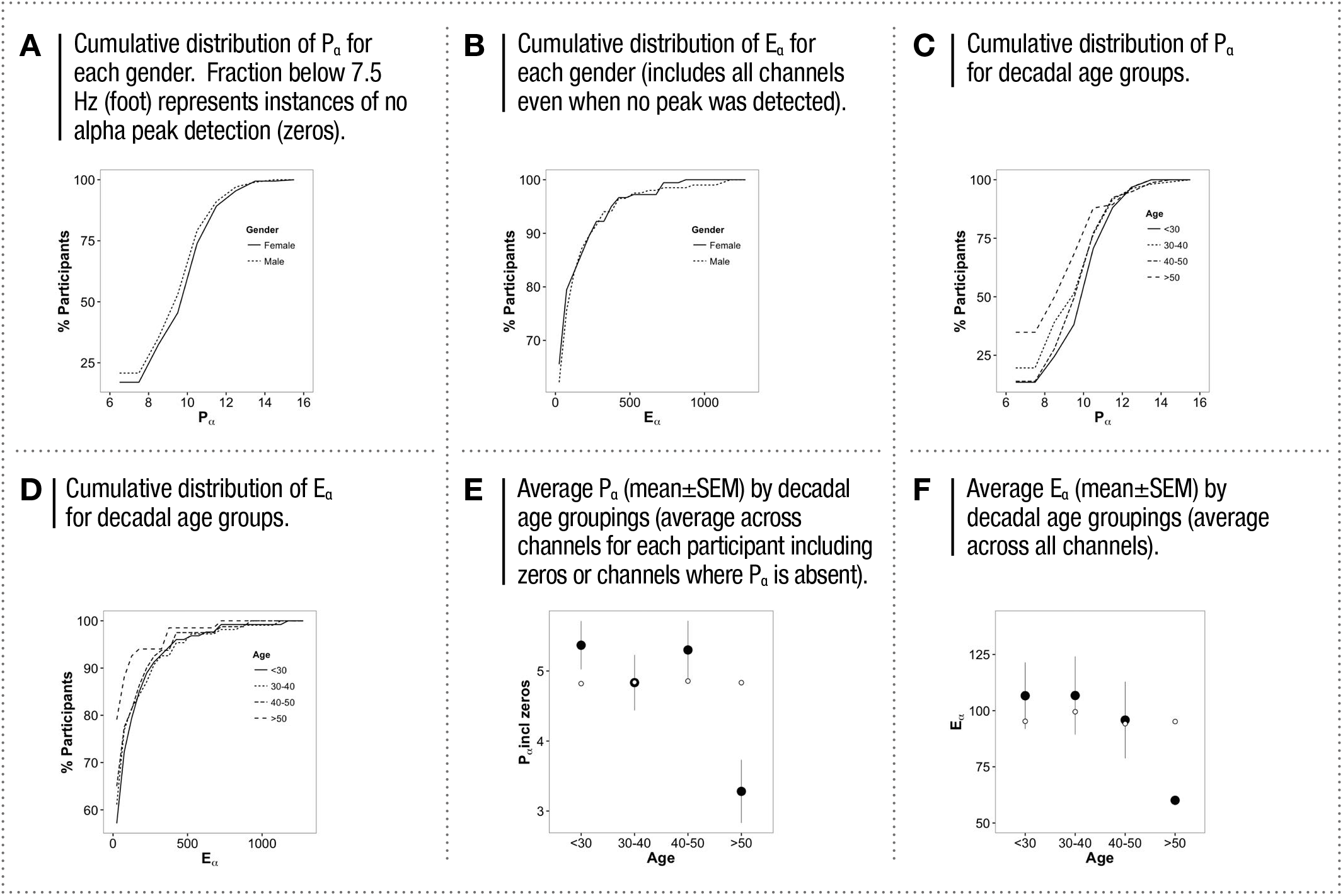
ALPHA OSCILLATION FEATURES BY AGE AND GENDER

We next looked at these parameters as a function of age (Fig. 5C, D). Previous studies have shown a decline with age, most significantly beyond the age of 45[27]. Here we found that there was no difference in *P_α_* (counted only when it was present) with age (Fig. 5C; ANOVA *F*=0.11, 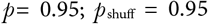, see methods for definition). The difference between the <30 group (10.49±0.11) and the >50 group (10.47±0.35) had a p-value of 0.95 (by t-test).

On the other hand, the fraction of channels with no detected alpha oscillation did differ across age groups (initial value of cumulative histogram in Fig. 5C.). Thus when the average *P_α_* was calculated including those channels with no detectable peak as zeros, a decline beyond age 50 relative to the under 30 was apparent (Fig. 5E; 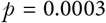 by t-test comparing <30 and >50; ANOVA for all groups *F*=4.73, *P*= 0.003; 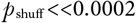).

In the case of *E_α_*, there was a weak decreasing trend with age. However, while the overall trend was not significant in our sample (ANOVA *F*=1.46, *P*= 0.22; 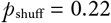), the over 50 age group was significantly lower than the under 30 group (Fig. 5D,F; 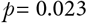 by t-test).

We also note that our sample was roughly equally split by age and gender groups across all income groups such that these results are not be likely to reflect other dimensions. Finally, we note that despite the significance of the difference after age 50, the magnitude of difference was not large enough to explain the very wide range of differences across our sample.

## THE ALPHA OSCILLATION AND MODERNIZATION

The fundamental difference between our sample population and those studied previously is that ours covered a large number of people with low income and limited exposure to modern life. We thus looked at the relationship between the peak and energy of the oscillation and modernization by dividing our sample into three groups of pre-modern, modern and transitioning (Fig. 6). We defined pre-modern as groups with only a primary education and below with no owned motorized vehicle or access to electricity and telecommunication and modern as those who were college educated or above with their own phone, vehicle and electricity. ‘Transitioning’ were all others who fell between these two extremes.

**Figure 6.**
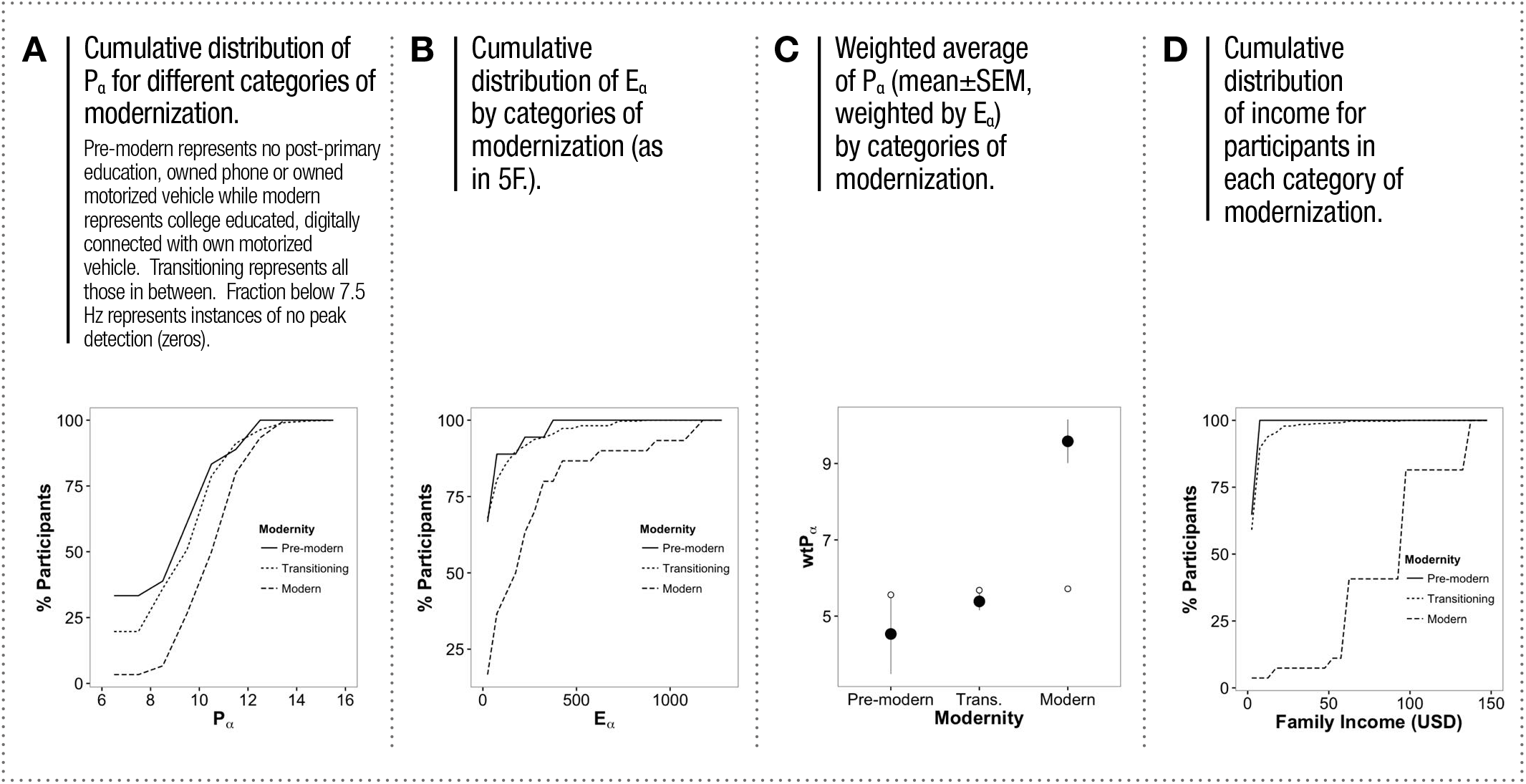
ALPHA OSCILLATION FEATURES BY MODERNIZATION

The average *P_α_* (when it was detected) was 10.28±0.31 for the pre-modern group, 10.43±0.11 for the transitioning group and 11.05 ±0.21 for the modern group. The pre-modern and transitioning were not significantly different from one another (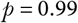 by KS test) but both together were significantly different from the modern group (Fig. 6A; 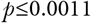 by KS test for both comparisons). The overall group comparisons however were not highly significant (ANOVA *F* = 2.55, 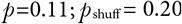).

On the other hand, the average *P_α_* across all channels, including those with no detected oscillations (i.e. *P_α_* =0), was significantly higher for the modern group (8.45±0.61) compared to both the pre-modern (3.91±0.95) and transitioning groups (4.56±0.21), (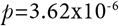 by KS test, ANOVA *F* = 14.84, 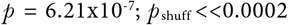). Whereas a peak was always detected in at least one channel in ~97% of those in the modern group, it was entirely absent in ~20.17% of the pre-modern and transitioning groups.

More dramatically, only 40% of the pre-modern and transitioning groups had *E_α_* beyond the non-oscillatory residual alpha energy level (defined as 2SD from the mean, here 39) while this was so for 93.3% of the modern group (Fig. 6B). *E_α_* was 64.44±21.5 for the pre-modern group and 81.49±7.2 for the transitioning group, which were not significantly different (p=0.9 KS-test). In contrast both were dramatically lower than the modern group, which had *E_α_* of 278.75 ±56.8 (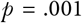 t-test, 3.2×10^-7^ KS-test, ANOVA *F*= 24, 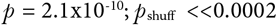).

The weighted *P_α_* (average *P_α_* weighted by *E_α_* and including channels with *P_α_* = 0 (i.e. no oscillation) was 4.53±1.05 for the pre-modern group, 5.40±0.23 for the transitioning group and 9.58±0.57 for the modern group (Fig. 6C).

Thus, taken together, the presence, peak and energy of the alpha oscillation is closely associated with access to features of modern living and its domination of the resting EEG may not simply be an intrinsic consequence of being born human but rather highly dependent on the context into which the person is born and raised.

The largest driving force with respect to access to the features of modernity is income, since all of the tools of modernity must largely be purchased. Fig. 6D shows the cumulative income distributions of each group in our sample. 65% of the pre-modern group had incomes under $5/day while the rest had incomes between $5 and $10/day. Incomes in the transitioning group were also largely under $10/day although 10% had incomes above $10. In contrast only 7% of the modern group had incomes below $30/day. This pattern suggests that there is a certain threshold of income that may be required to have sufficient access to the tools of modernity and thereby drive a rapid change in alpha oscillations.

Taken together these results demonstrate a vast divergence in the presence of the alpha oscillation in the resting state across humanity that is markedly stronger in those beyond a certain threshold of access to the tools and structures of modern civilization.

## DISCUSSION

Here we have shown that the alpha oscillation, thought to be a fundamentally ubiquitous and dominant feature of brain activity, widely diverges across humanity in its presence in the EEG. Furthermore, the energy of this oscillation has dramatic variance, distributing across our sample with a 1000x range with no centralizing mean. This raises a very fundamental question about how one might define the ‘average’ human brain in a dynamical context and the surprising answer that it perhaps does not exist. Unlike other organs, such as the heart, where dynamical markers of function might be easily used to create benchmarks that guide diagnosis of normal and disease states, using such a marker for the brain poses unique problems. These results suggest that ‘normal’ may have to be related to the specific context of human experience.

Our most significant finding, however, is that the ubiquitous emergence of this feature is related to modern experience. Considering that the alpha oscillation has several cognitive implications, this suggests that modern experience may change the cognitive capacity of the human brain quite profoundly. This is even more poignant when one considers that while modern civilization has emerged only over the past 200 years, it has not done so uniformly across the globe. Furthermore, with incomes having diverged so dramatically over the past century structuring as a long tailed distribution, it is only a minority of humans that have the ability to afford all the tools of modern living. Indeed, only 15-20% of humanity today would be considered ‘modern’ by our definition. With the average weighted *P_α_* across channels shifting nonlinearly and clear evidence for the functional cognitive associations with the alpha oscillations, this suggests that we might be diverging into populations of distinct and divergent modes of cognitive functioning. It is also of significance that the alpha oscillation is not typically present in young children, emerging only after age five and growing over childhood to reach maturity in later years [27]. This suggests that it is largely an experience dependent phenomenon of the brain. This would have serious implications for how we approach education and social policy across different populations.

Parsing the effects of different aspects of modernization such as education and technology will therefore be of considerable interest, as also a deeper understanding of the implications of this divergence for various types of cognitive abilities.

## ACKNOWLEDGEMENTS

A very special thank you to Sathish S, Aravind and Govind for managing the field EEG recordings and surveys, as well as S. Ravi Shankar who enabled smooth coordination of the process. We also thank the various staff at Madura Microfinance who facilitated access to the villages and recruitment of participants and SciSphere, India for providing us survey tools, access to demographic and economic data for location selection and data support. We also thank Dietmar Plenz and Christian Meisl for useful discussion.

